# Accurate sequence-to-affinity models for SH2 domains from multi-round peptide binding assays coupled with free-energy regression

**DOI:** 10.1101/2024.12.23.630085

**Authors:** Dejan Gagoski, H. Tomas Rube, Chaitanya Rastogi, Lucas A. N. Melo, Xiaoting Li, Rashmi Voleti, Neel H. Shah, Harmen J. Bussemaker

## Abstract

Short linear peptide motifs play important roles in cell signaling. They can act as modification sites for enzymes and as recognition sites for peptide binding domains. SH2 domains bind specifically to tyrosine-phosphorylated proteins, with the affinity of the interaction depending strongly on the flanking sequence. Quantifying this sequence specificity is critical for deciphering phosphotyrosine-dependent signaling networks. In recent years, protein display technologies and deep sequencing have allowed researchers to profile SH2 domain binding across thousands of candidate ligands. Here, we present a concerted experimental and computational strategy that improves the predictive power of SH2 specificity profiling. Through multi-round affinity selection and deep sequencing with large randomized phosphopeptide libraries, we produce suitable data to train an additive binding free energy model that covers the full theoretical ligand sequence space. Our models can be used to predict signaling network connectivity and the impact of missense variants in phosphoproteins on SH2 binding.

## INTRODUCTION

Many protein-protein interactions in the cell occur between a peptide-recognition domain and a short linear peptide sequence, both of which are typically embedded within larger proteins^1^. These peptide sequences, often referred to as short linear motifs (SLiMs), can be bound by a variety of peptide recognition domains (PRD), with a binding energy that can depend strongly on the amino-acid sequence of the peptide ligand. SLiMs play important roles in the formation of protein complexes and the regulation of signaling cascades. Paralogous individual PRDs from the same family can have distinct SLiM binding preferences indicative of functional specialization, despite their overall homology. SLiMs often contain residues that can be post-translationally modified, providing a means for positive or negative regulation of their interactions. For example, Src-homology 2 (SH2) domains specifically bind to SLiMs centered on phosphorylated tyrosines, and as such, they mediate regulated protein-protein interactions in response to tyrosine kinase activity.

Notably, mutations within SLiMs can either weaken or strengthen the protein-protein interactions they mediate, allowing for rapid evolution of new signaling networks as well as the emergence of pathogenic molecular processes^2–4^. Our ability to predict the wiring and rewiring of SLiM-based interaction networks could enhance fundamental studies on cell signaling, shed light on the evolution of signaling pathways, reveal mechanisms of pathogenicity for uncharacterized mutations, and ultimately inspire novel therapeutic strategies.

Over the past two decades, many platforms for probing SLiM-based interaction networks have been developed^5^. These approaches rely on a range of different techniques, including synthetic peptide arrays^6–9^, two-hybrid assays^10^, affinity purification coupled with mass spectrometry^11^, and display technologies^12,13^. What these platforms have in common is that they aim to rapidly assess the binding of a peptide-recognition domain against a large library of peptides or proteins, towards the goal of identifying optimal binders or obtaining a description of sequence preference (typically in the form of a consensus motif or a sequence logo). The use of increasingly large and affordable DNA-encoded protein/peptide display libraries coupled with deep sequencing has dramatically increased the scale and throughput of these experiments^12,14–19^.

Datasets generated using such high-throughput methods are well-suited for the development of *quantitative* models of sequence specificity, the simplest of which is the widely-used position-specific scoring matrix (PSSM). These matrices are typically derived by first aligning the sequences of all experimentally-determined “binders” around a common register (e.g., a phosphorylated amino acid), and then determining the frequency of each amino acid at each position surrounding that reference point. The weights are sometimes normalized to the amino acid frequencies in the naïve input library that includes both bound and unbound sequences. The resulting matrix of amino-acid identity by residue position provides a simple way to score and rank sequences by relative enrichment, irrespective of their presence in the original dataset. In cases where entire families of peptide-recognition domains have been experimentally characterized, these matrices provide a powerful way to differentiate binding specificities among closely-related proteins, as recently exemplified by landmark studies on the human “kinome” ^20,21^. However, there are also drawbacks to the use of position-specific scoring matrices, including the requirement to pre-define a binding register and an inability to account for interdependencies between residue positions or non-specific binding.

Large-scale SLiM recognition datasets also lend themselves well to statistical learning approaches that treat prediction of peptide-protein interactions as a binary classification problem and go beyond the PSSM representation. These include support vector machines^22,23^ and random forest classifiers^24,25^. Deep learning approaches that combine sequence and structural data have also been explored^26^, as has the general sequence-to-structure predictor AlphaFold 2^27,28^. For SH2 domains in particular, peptide array data have been used in combination with non-linear support vector machines^29^ and deep learning to classify peptide-SH2 domain interactions as strong or weak (Tinti et al., 2013). In both cases, a goal was to capture dependencies between amino acid residue positions within the bound peptide. However, these approaches tend to be hampered by the oversampling of positive interactions^30^.

Datasets generated more recently using genetically-encoded libraries and deep sequencing hold particular promise for machine learning approaches, as the high data volume and broad sampling of sequence space relative to other methods may allow for better model training. Recently, several SH2 domains were analyzed using bacterial display of genetically-encoded peptide libraries, enzymatic phosphorylation of the displayed peptides, affinity-based selection, and deep sequencing^31–33^. These studies employed a variety of library formats, including scanning mutagenesis libraries derived from individual peptides (<10^3^ sequences), phosphoproteome-derived peptide libraries (10^3^-10^4^ sequences), and degenerate libraries (10^6^-10^7^ sequences).

Here, we report a coordinated experimental and computational strategy for analyzing sequence recognition by PRDs (**Figure 1**) that improves upon these pioneering studies. Our approach employs ProBound, a statistical learning method originally developed by our group to accurately model protein-DNA interactions^34^. We illustrate how model quality can be improved by using data from multiple rounds of selection on highly degenerate libraries. Focusing on SH2 domains as a case study, we demonstrate that unbiased selection and sequencing datasets generated using highly degenerate random libraries can be used to build models that accurately predict binding affinity for any ligand sequence in the theorical space covered by the library. Our strategy should be readily adaptable to other peptide recognition domains.

**Figure 1:**
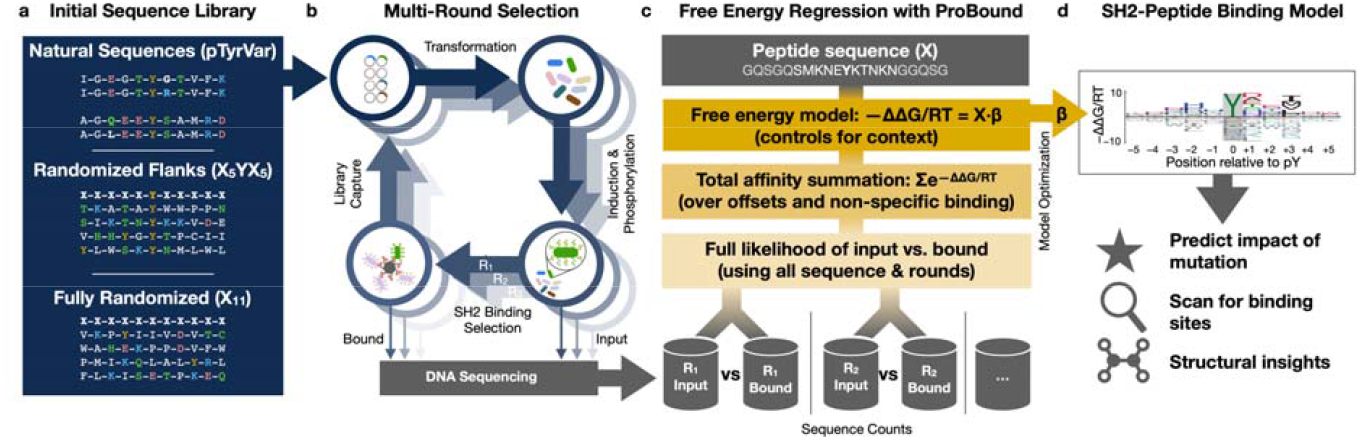
Overview of concerted experimental and computational strategy for generating SH2-peptide binding free energy models. (**a**) Design of peptide-display libraries. (**b**) Schematic showing how a randomized bacterial display library underwent repeated bead-based affinity selection for SH2 binding. In each selection round, the library was sequenced before and after selection. (**c**) Overview of the regression framework used to learn energetic binding models from the sequencing data. For each possible binding site, the energy received independent additive contributions from the residues flanking the phosphorylated tyrosine, thus controlling for the binding-site context wherein the residues reside. These energy contributions were estimated using maximum likelihood estimation, where the likelihood of the observed sequence counts was evaluated by first computing the total affinity for each observed sequence (controlling for multiple possible binding offsets and non-specific binding) and then computing the binomial likelihood for each round, assuming linear section. (**d**) Sequence logo displaying the inferred energy contributions as letters whose height reflects the magnitude of the contributions, relative to the mean for each position.

## RESULTS

### Robust inference of SH2-peptide binding free energy models using ProBound

A recent study involving some of the authors of this work^33^ used bacterial surface display of plasmid-encoded peptides containing a central phosphorylated tyrosine and deep sequencing to assay the target specificity of the SH2 domain from c-Src kinase. Two distinct library designs were used: one (“pTyrVar”) based on ∼10^4^ peptides containing a phosphorylated tyrosine and occurring in the human population, the other a synthetic random library (“X_5_YX_5_”) with a fixed tyrosine between two fully degenerate five-amino acid flanks with a theoretical diversity of ∼10^13^ and an actual diversity of ∼10^6^ (**Figure 1a**). Importantly, this study relied on position-specific maps of relative enrichment, obtained by comparing amino-acid frequencies directly before and after affinity-based enrichment, to provide a simple and intuitive way to summarize the sequence preferences of the SH2 domain and predict relative binding affinities of unseen peptides^33^.

We hypothesized that using relative enrichment as a proxy for the true binding free energy differences (ΔΔG/RT) associated with amino-acid substitutions in the peptide could be suboptimal from a quantitative point of view: Relative enrichment may depend on library design, whereas binding free energy differences, being an intrinsic property of the SH2-peptide interface, should not. To test this, we first compared peptide enrichments from the pTyrVar and X_5_YX_5_ libraries upon selection by the c-Src SH2 domain (**Figure 1b**). We observe that a larger fraction of sequences showed strong enrichment in the pTyrVar library over the X_5_YX_5_ library (**Figure 2a**), resulting in apparently stronger amino acid preferences with the pTyrVar library even though the screens were conducted under identical conditions (**Figure 2b-c**).

**Figure 2:**
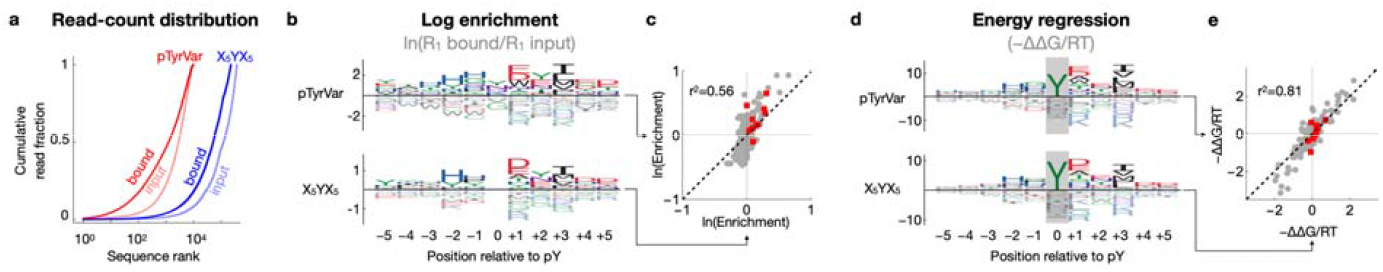
Comparison of amino-acid enrichment analysis and free-energy regression. (**a**) Distribution of read counts (after down-sampling to 500,000 reads) for sequences in the pTyrVar and X_5_YX_5_ libraries, respectively, each before and after one round of affinity selection with the c-Src SH2 domain. (**b**) Amino-acid log-enrichment due to affinity selection for c-Src SH2, displayed as sequence logos, for the designed pTyrVar and random X_5_YX_5_ library, respectively. (**c**) Direct comparison of log-enrichment parameters between the two library designs. Red points again indicate tyrosine. (**d**) Inferred free-energy contributions (ΔΔG/RT) at different positions within the c-Src SH2 binding interface, displayed as sequence logos. Gray rectangles indicate position where the model was constrained to recognize (phospho)tyrosine. (**e**) Direct comparison of ΔΔG/RT parameters between the two library designs.

Our next goal was to assess whether ProBound^34^ is also capable of building accurate sequence-to-affinity models from high-throughput protein-peptide binding data (**Figure 1c**). To this end, we configured ProBound to learn a free-energy matrix that encodes how the SH2 domain interacts with an 11-amino-acid subsequence (**Figure 1d**). To focus this matrix on sequence-specific SH2 binding, the central column was constrained to recognize tyrosine. The non-central columns were learned using maximum likelihood estimation over the input and binding-selected libraries. Specifically, the selection of each sequence was modeled by first computing the total sequence-specific binding affinity (scoring and summing over all binding offsets, thus controlling for non-central tyrosines) and then adding a non-specific term to capture background selection and simple sequence biases (see **Methods**).

We found that the ΔΔG/RT parameters in the resulting models were far more consistent between the two library designs (**Figure 2d-e;** *r*^*2*^ = 0.81) than the corresponding log-enrichments (**Figure 2b-c**; *r*^2^ = 0.56); this is likely due to the fact that ProBound controls for sequence context and the effect of non-specific binding, both of which are library dependent, when estimating the energetic effect of amino-acid substitutions. Moreover, the log-enrichment analysis indicated selection for tyrosine at non-central positions (**Figure 2b**), a likely artifact due to the fact that the libraries are enzymatically phosphorylated and contain non-central phosphotyrosine residues, which results in SH2 binding at non-central offsets; by contrast, the ProBound models, which account for all offsets, do not show disproportionate enrichment for tyrosine at non-central positions (**Figure 2d**). Together, these results demonstrate that ProBound has superior robustness with respect to library design.

### Feasibility of using fully random libraries over multiple successive rounds of selection

The library designs used so far exploited prior knowledge about the SH2 domains, specifically, their strong requirement for a phosphorylated central tyrosine residue at the binding interface ^35,36^. Such a biased design, however, may be undesirable or unfeasible when a similar experimental strategy is to be used to characterize other peptide binding domains. We therefore repeated our peptide binding assay using a new library (“X_11_”) in which 11 consecutive residue positions are fully randomized; we otherwise followed the protocol of ^33^, including phosphorylation of the bacterial display library before c-Src SH2 domain binding. The results are shown in **Figure 3a**. Using ProBound to model the evolution of the X_11_ library over a single round (R_1_) of selection yielded binding free energy parameters much less consistent with those we obtained using the X_5_YX_5_ library (**Figure 3b**; *r*^*2*^=0.63). Since X_11_ is expected to be more dominated by weak binders compared to the other two libraries, it was perhaps not surprising that the R_1_ data for X_11_ do not contain adequate signal for successful binding model inference. To address this, we developed a multi-round selection strategy that aims to maximize recovery of the bound fraction of the library in each round (see **Methods**). In addition to the bound fraction, the input was also sequenced as a control for each round, and ProBound was configured to jointly learn from all input-output pairs across all rounds. Using this protocol to generate additional data, we found that three rounds of selection with the X_11_ library were necessary and sufficient to obtain adequate signal for building high-quality sequence-to-affinity models for the c-Src SH2 domain using ProBound (**Figure 3c**) that agreed well with the model built from R_1_ of the X_5_YX_5_ library (**Figure 3d**; *r*^*2*^ = 0.82).

**Figure 3:**
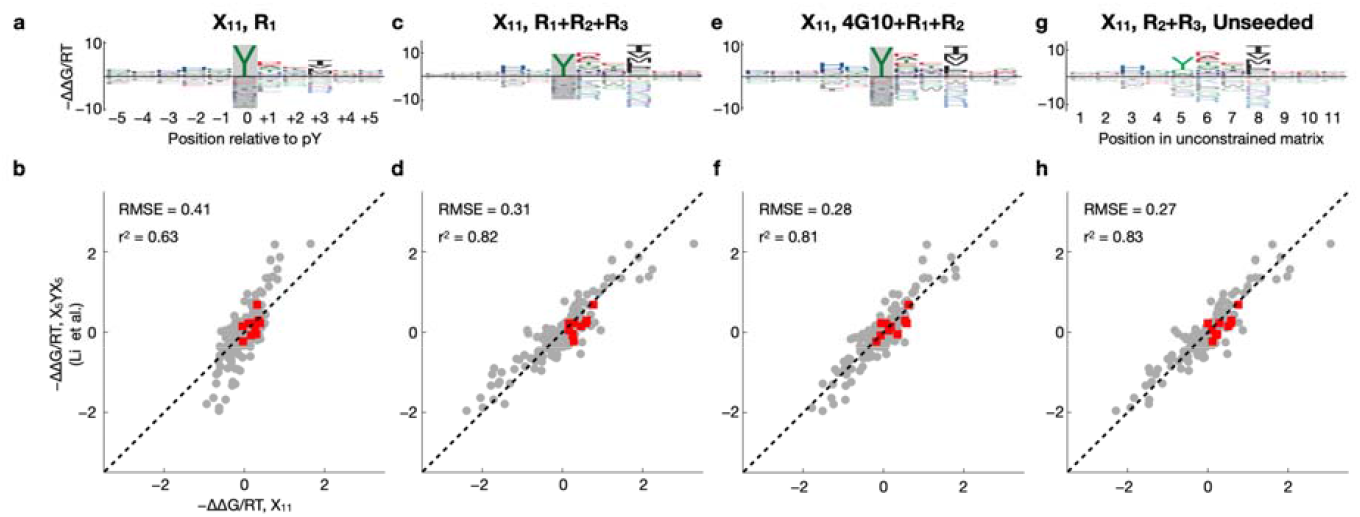
Multi-round profiling of c-Src SH2 using the naïve and pre-enriched X_11_ libraries. (**a**) Binding model learned using one selection round and starting with the naïve X_11_ library. (**b**) Scatter plot comparing the model coefficients shown in panel (a) to the coefficients of the X_5_YX_5_ model shown in Fig. 2d. Red points indicate tyrosine. (**c-d**) Same as (a-b) but showing a model that was trained on data from three selection rounds. (**e-f**) Same as (a-b) but showing a model that was trained on an experiment where the input library was pre-selected using the 4G10 antibody, followed by two rounds of c-Src SH2 binding selection. (**g-h**) Same as (a-b) but showing a model that was trained on data from the second and third selection rounds and that was not constrained to recognize tyrosine at the central position.

A disadvantage of using the unbiased X_11_ library to profile the binding preferences of SH2 domains is that many sequences in the input library will not contain any tyrosine residues (**Figure S1**). Moreover, depending on its flanking sequence, not every Tyr will be as efficiently phosphorylated by the kinases used for enzymatic phosphorylation. Because specific recognition of pTyr is a major contributor to SH2 binding affinity, we performed target-agnostic pre-selection on the X_11_ library using a biotinylated anti-phosphotyrosine antibody (4G10) as described in ^33^. We found that two subsequent rounds of c-Src SH2 domain selection were sufficient in this case to obtain a high-quality binding model (**Figure 3e**) that agrees well with the X_5_YX_5_ model (**Figure 3f**; *r*^*2*^ = 0.81).

An advantage of the X_11_ library its universality: Even proteins with completely unknown sequence preferences could in principle be characterized in an unbiased manner given enough selection. We therefore asked if the c-Src SH2 domain could be characterized without using the prior knowledge that having a tyrosine at the central position is required for binding. Reconfiguring ProBound to learn a completely unconstrained free energy matrix from the R_2_ and R_3_ libraries produced a binding model (**Figure 3g-h**) whose ΔΔG/RT parameters again agreed well with those of the centrally constrained model built from R_1_ of the X_5_YX_5_ library (*r*^2^ = 0.83).

### Quantifying differences in flanking sequence preference among paralogous SH2 domains

To test the ability of our approach to resolve differences in binding specificity between paralogs, we performed two-round selections of the X_5_YX_5_ library against the SH2 domains of two closely-related kinases from the Src subfamily (c-Src and Fyn) and of one distantly-related SH2 domain from the adapter protein Grb2. For the Grb2 SH2 domain, we observed a strong preference for an asparagine (N) residue at position +2 relative to the pTyr (**Figure 4a**), consistent with previous findings^37^. Notably, for Grb2, the relative peptide binding affinities predicted by our model for peptides with and without an N_+2_ were separated by 2-3 orders of magnitude (**Figure S2**). This is consistent with previous measurements of Grb2 SH2 affinity, which indicate that substitution of the N_+2_ residue disrupts binding affinity by at least 100-fold^38^.

**Figure 4:**
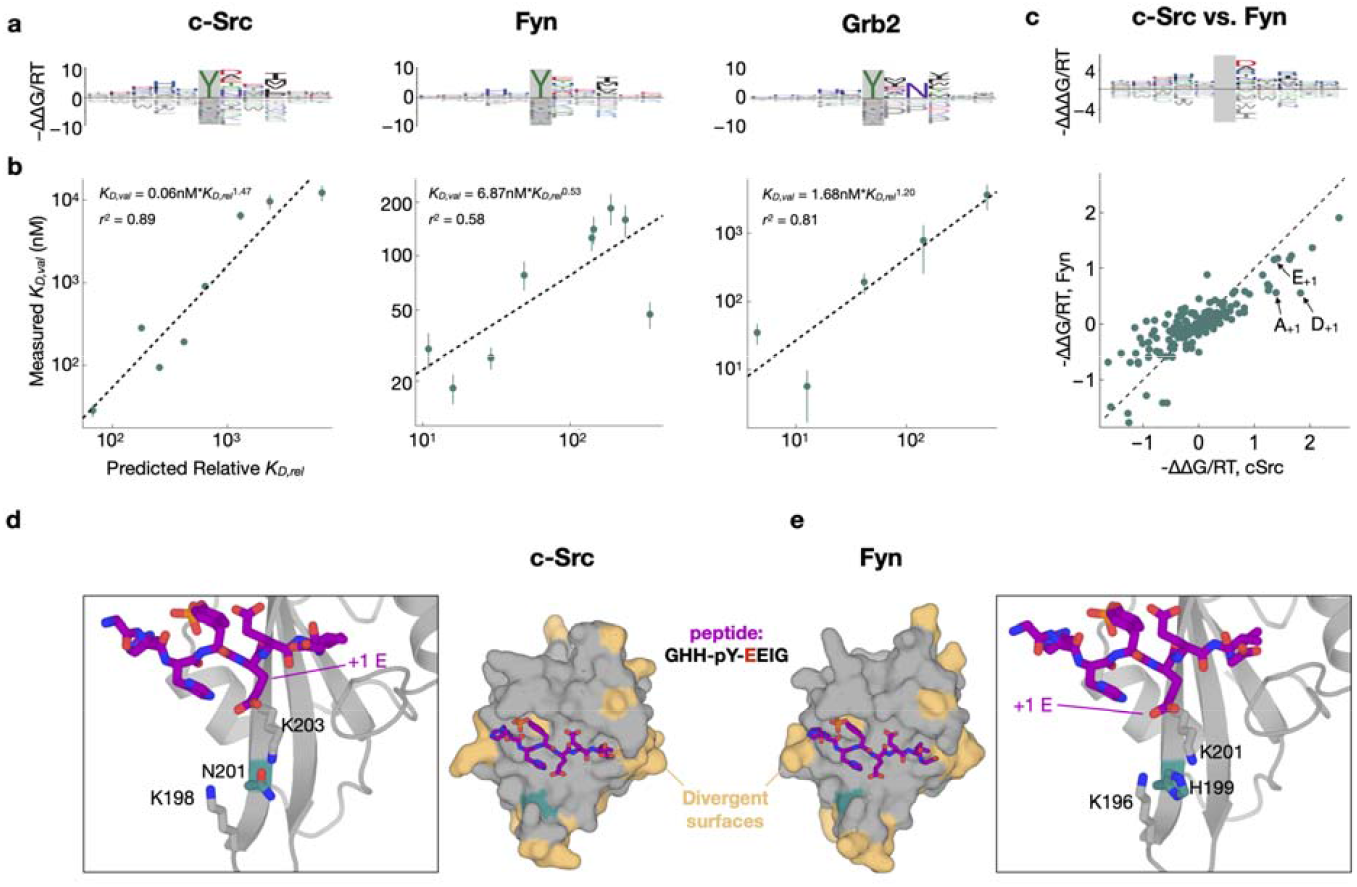
Flanking specificity of the c-Src, Grb2 and Fyn SH2 domains. (**a**) Energy logos for the c-Src SH2, Grb2 SH2 and Fyn SH2 binding models highlighted by arrows in (a). (**b**) Scatter plots comparing the predictions from the binding models in (a) with competitive fluorescence polarization measurements. Vertical bars indicate standard error. Dashed black lines (and accompanying model expressions and r^2^ values) indicate linear regression fits to the log-transformed K_D_-values. (**c**) Comparison of the c-Src and Fyn binding models from (a) using an energy logo (top, showing the difference between the model coefficients) and a scatter plot (bottom). (**e-f**) AlphaFold 3 models of the c-Src and Fyn SH2 domains (shown as surfaces in the central panels) bound to a high-affinity phospho-peptide (GHH-pY-EEIG, shown as purple sticks). Residues on the SH2 domains colored in beige are sites where c-Src and Fyn diverge. A key divergent site (N201 in c-Src and H199 in Fyn) is shown in teal. The zoom-in panels highlight key residues in a cationic pocket on the SH2 domain that interacts with the +1 residue on the peptide ligand.

As a more stringent and direct experimental test of the predictive capability of our sequence-to-affinity models, we conducted low-throughput competition fluorescence polarization assays for a panel of individually synthesized phosphopeptides covering almost two orders of magnitude of predicted binding affinity (see **Methods**). The measured ln(K_D_) values for each of the three SH2 domains showed good to excellent agreement with the corresponding ΔΔG/RT values predicted using our ProBound models (**Figure 4b; Table S3;** *r*^2^ values ranging from 0.58 to 0.89).

When comparing the c-Src and Fyn models, we noted a few reproducible differences in specificity. For example, whereas both domains preferred a glutamic acid at the +1 position relative to the tyrosine (E_+1_), c-Src had a distinctive preference for aspartic acid (D_+1_) and alanine (A_+1_) relative to Fyn (**Figure 4c**). To rationalize these differences, we used AlphaFold 3^39^ to generate models of the c-Src and Fyn SH2 domains bound to a predicted high-affinity phosphopeptide bearing a E at the +1 position. The c-Src and Fyn SH2 domains have 66% sequence identity, with very few divergent positions within the phosphopeptide binding pocket (**Figure 4d-e**). One such residue is N201 in c-Src, which corresponds to H199 in Fyn. This residue sits at the center of a positively charged pocket that coordinates E_+1_ (**Figure 4d-e**). Our structural models and affinity models suggest that alteration of this central residue from an obligate neutral amino acid in c-Src (N) to a potentially charged amino acid in Fyn (H), changes both the steric and electrostatic surface potential in this region, thereby altering amino acid preferences at peptide position +1.

Finally, in a more comprehensive test of the feasibility of using a target-agnostic library approach, we also performed three rounds of selection of X_11_ against the same set of SH2 domains. Apart from one exception, the binding models inferred from the data we generated (**Figure S3**) cluster by SH2 domain identity (**Figure S4; Tables S1-2**). This indicates that ProBound is appropriately controlling for the significant differences between the X_11_ or X_5_YX_5_ library, and whether 4G10 pre-selection was used or not.

### Affinity models enable the discovery of putative novel interactions

Having validated their accuracy experimentally, we sought to investigate whether our SH2 domain affinity models could be used to assign relative affinities to known phosphorylation sites in the human proteome, with the goal of identifying putative interaction partners not previously reported in the literature. First, we generated bacterial display datasets and affinity models for three additional SH2 domains from Src-family kinases Lyn, Blk, and Yes (**Figure 5a**). For validation of the affinity model for Lyn, we selected a subset of phosphopeptides that were likely to fall within the dynamic range of the competitive fluorescence polarization assay (see **Methods** for details). We selected 9 phosphorylation sites that should bind to the Lyn SH2 domain with moderate to high affinity, including 3 previously reported interactors (PLCγ2 pY753, CSFR pY699, and SHP1 pY564) and 6 sites not previously reported to bind to Lyn (CD3ζ pY83, SLAP-130 pY771 and Y595, TRAF3IP3 pY322, CD84 pY296, and SIGLEC5 pY544) in the STRING database^40^. We measured binding affinities to the corresponding phosphopeptides by fluorescence polarization. The Lyn SH2 domain showed a good correlation between measured binding affinities and relative affinities predicted by the model (**Figure 5b**; *r*^2^ = 0.82), indicating that this should be a reliable strategy to identify putative SH2 interactors across the proteome.

**Figure 5:**
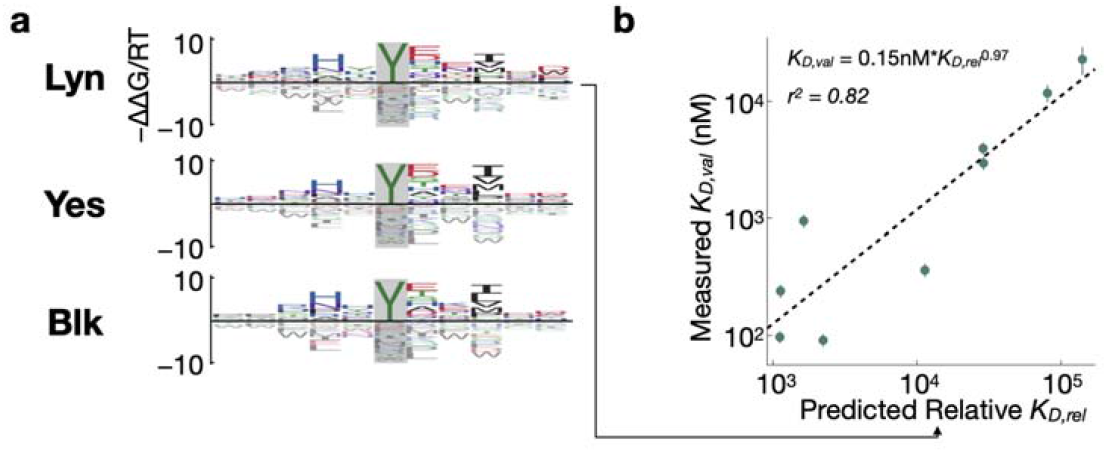
Flanking specificity for the Lyn, Yes and Blk SH2 domains. (**a**) Energy logos showing binding models for Lyn, Yes, and Blk. The models were trained on two-round experiments using the X_5_YX_5_ starting library. (**b**) Scatter plots comparing model predictions and validation measurements for the Lyn SH2 domain, shown as in Figure 4b.

We next used our models to predict relative affinities for all of the human tyrosine phosphorylation sites documented in the PhosphoSitePlus database^41^. Among the proteins containing these phosphorylation sites, we first filtered out those that do not co-express with the given SH2-containing kinase according to the STRING database^40^. The lists of predicted high-affinity binders for each SH2 domain (**Table S4**) show that majority of the phosphorylation sites have low binding scores, indicating the preference of the SH2 domains towards a relatively small number of phosphorylation sites. It is noteworthy that none of the phosphorylated sites reach the maximal theoretical binding score, suggesting that few, if any, phosphosites in the proteome have evolved to be bound by their cognate SH2 domains at the highest possible binding affinity.

### Affinity models correctly predict effects of single-amino-acid substitutions in SH2 ligands

The fact that SH2 domains display position-specific sequence preferences means that single-amino-acid substitutions flanking the phosphorylated tyrosine could have a significant effect on SH2 binding affinity, and this should be captured by our Probound models. To test this, we produced several pairs of phosphorylated peptides derived from human proteins, with one peptide in each pair a reported wildtype sequence and the other containing a single-amino-acid mutation reported to be a natural human variant^41^ (**Table S5**). We experimentally measured binding affinities for these peptides against the c-Src and Fyn SH2 domains and also predicted their relative affinities using the corresponding Probound models. For both SH2 domains, the change in measured binding affinity due to the mutation had the same directionality as predicted by the affinity models (**Figure S5**).

We next used the Probound models to score all sequences in the PTMVar database of human phosphorylation site variants^41^, to predict if the reported variants enhance or diminish binding relative to the wild type sequence^41^ (**Table S6**). Notably, many of the variants in this database are derived from patients with specific disease phenotypes, and these mutations may contribute to rewired pathogenic signaling events. Filtering for proteins that co-express with a given SH2 domain and removing variants with substitution of the phosphorylated tyrosine itself, yielded the list of predicted variant effects summarized in **Figure 6**. Several of these variants could substantially alter cell signaling in a biologically meaningful way. For example, the c-Src SH2 domain exhibits loss of affinity for Vav2 at the Y142 phosphorylation site in the presence of a lung cancer variant, R143L. c-Src is directly involved in tyrosine phosphorylation at this same site in Vav2^42^ and both proteins co-localize with EGFR, a known driver of lung cancer. Our models also predict that the Fyn SH2 domain has an increased affinity for the Y492 phosphorylation site on ZAP-70 in the presence of the A495V variant. Both ZAP-70 and Fyn play a key role in mediating T cell activation downstream of the T cell antigen receptor, and Y492 and adjacent Y493 on ZAP-70 both are critical phospho-regulatory sites in this pathway^43^. For the Lyn SH2 domain, we predict a loss of affinity towards the HS-1 Y360 phosphorylation site when the G361K variant is present. This is likely to have implications in the context of the Lyn/HS-1 signaling axis, which is of therapeutic interest for Chronic Lymphocytic Leukemia^44^. Finally, for the Grb2 SH2 domain, we observed a strong enhancement in binding towards the Y35 site of ARPC1B with the K37N variant. Given the role of ARPC1B in actin cytoskeleton scaffolding, this neo-interaction with Grb2, a ubiquitous adaptor protein, could alter the recruitment of other signaling proteins to the cytoskeleton during important cellular events, such as T cell recognition of antigen presenting cells, where both ARPC1B and Grb2 have already been functionally implicated^45^.

**Figure 6:**
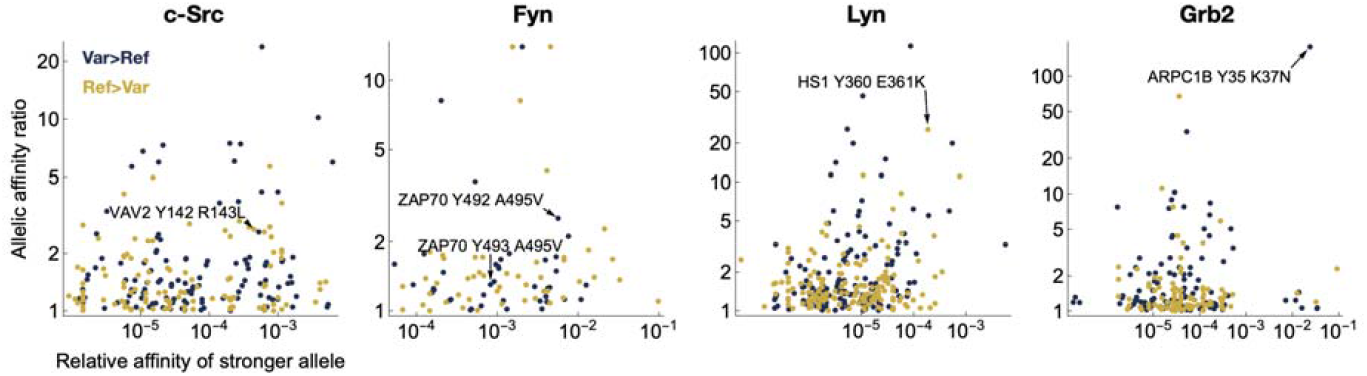
Distribution of the predicted quantitative impact of missense variants in SH2 binding sites in the human proteome. Scatterplot of allelic effect of missense variation in SH2 binding sites documented in the PTMVar database of human phosphorylation site variants ^1^, colored by the direction of the effect. The x-value corresponds to the greater of the predicted affinities of the two alleles; the y-value corresponds to the ratio of predicted affinities between the two alleles.

## DISCUSSION

In this study, we have demonstrated how bacterial display of peptide libraries, multi-round affinity-based selection, and deep sequencing can be combined with principled regression of binding free energy parameters to obtain robust models of peptide binding specificity. Our free-energy regression approach has several advantages over simpler enrichment-based strategies. First, when estimating the energetic impact of an amino acid substitution, the confounding impact of the other residues in the binding site is controlled for by predicting the affinity of the full binding interface, rather than simply computing the amino-acid enrichment at each position independently. Similarly, the confounding impact of possible secondary binding sites is controlled for by considering all possible offsets and predicting the total binding affinity. Finally, data were synthesized by fitting a single binding model that best explains the full multi-round experiment. Together, these improved computational methods enable an unbiased, target-agnostic approach using highly-complex random libraries.

Applying our methodology to various peptide binding domains from the SH2 family, we found that the predictive accuracy of the models depends strongly on the identity of the SH2 paralog, as well as on the design of the selection assay used to generate the training data. The presence of a post-translationally modified phosphotyrosine in the center of the bound peptide, which contributes strongly and specifically to the binding affinity for all SH2 domains, may have contributed to the success of our current approach.

We found that it is important to objectively quantify the accuracy of each model by obtaining low-throughput measurements of binding constants for a small set of synthesized peptides and directly comparing these to the model’s predictions of relative affinity. However, once our sequence-to-affinity model for a particular SH2 domain passes this stringent quality assessment step, it is likely to give reliable predictions across the full theoretical space covered by the library on which it was trained. As such, it is a valuable tool for predicting putative new targets of the SH2 domain in the cell’s protein-protein interaction network, or for predicting the quantitative effect of allelic variation in the ligand sequence on SH2 binding affinity

The human genome encodes a great variety of peptide binding domains^1^. To make progress in understanding the structure and function of the protein interaction networks defined by these domains, it will be essential to construct high-quality sequence-to-affinity models for each of them, and move beyond the binary classification in terms of targets and non-targets that characterizes currently available resources^46^. The application of peptide display coupled with deep sequencing should make it feasible to generate high-throughput training data, and perform unbiased analyses similar to what we did here for SH2 domains, across the many other biologically important families of peptide recognition domains.

## MATERIALS AND METHODS

### Cloning, expression, and purification of SH2 domains

The human SH2 domains from Fyn (142-248), Lyn (121-228), Blk (116-222) and Yes (150-257) kinases were PCR-amplified from plasmids encoding the full-length proteins and cloned using the NEBuilder® HiFi DNA Assembly Master Mix (NEB) in a pET plasmid backbone that is PCR-amplified with the primers pETXbaI_for and pETXhoI_rev (these and all other primers are defined in **Table S7**) from the pET-His6-SUMO-Grb2(SH2) as reported before^33^, thus adding a N-terminal His_6_-SUMO tag and a C-terminal AviTag® for biotinylation. For expression and parallel biotinylating of the SH2 domain product *E. coli* C43(DE3) cells carrying a BirA encoding helper plasmid were transformed with the respective SH2 domain expression construct, and plated to grow over-night at 37 °C on a streptomycin and kanamycin (100 µg/mL both) LB-agar plate. The colonies were collected in 10 mL of TB-medium and used to inoculate 1L of TB medium and grown at 37 °C and 215 rpm. Upon reaching an OD_600_ of 0.5, the expression of the SH2 domains was induced by adding IPTG up to 0.5 mM biotin up to 250 μM, and the expressing culture was incubated at 18 °C for 16-18 hours. After removing the medium through centrifugation, the cell pellets were snap-frozen in liquid N_2_ and stored at -80 °C until purification. To purify the biotinylated SH2 domains the cell pellets were resuspended in lysis buffer (50 mM Tris, pH 7.5, 300 mM NaCl, 20 mM imidazole, 10% glycerol), supplementing them with 2 mM β-mercaptoethanol and 1x protease inhibitor cocktail before lysis by sonication (Fisherbrand Sonic Dismembrator). The soluble fraction separated after centrifugation at 33,000 g was run on HisTrap HP column (Cytiva), then washed with 10 CV of the lysis buffer followed by 10 CV of wash buffer (50 mM Tris, pH 7.5, 50 mM NaCl, 10 mM imidazole, 10% glycerol). Next, the protein was eluted with elution buffer (50 mM Tris, pH 7.5, 50 mM NaCl, 250 mM imidazole, 10% glycerol) directly onto a 5 mL HiTrap Q HP anion exchange column (Cytiva). The biotinylated SH2 domain was then eluted from the anion exchange column with an NaCl gradient (50 mM Tris, pH 7.5, 0.05-1 M NaCl, 1 mM TCEP-HCl) and the His_6_-SUMO tag was cleaved using 0.05 mg/mL Ulp1 protease. The biotinylated SH2 domain was then collected in a flowthrough after running the cleavage reaction over a HisTrap HP column. Finally, the biotinylated SH2 domain in the flowthrough was run on a HiLoad 16/600 Superdex 75 pg size exclusion column (Cytiva) in storage buffer (20mM HEPES, pH 7.5, 150mM NaCl, 10% glycerol). The purified biotinylated SH2 domains were aliquoted, snap-frozen in liquid N_2_ and stored at –80 °C.

### Expression and purification of kinases

The expression constructs for the four kinase domains EPHB1 (pET23a-H6-TEV-EPHB1; AddGene: 79694), c-Src (pET23a-H6-TEV-cSrc; AddGene: 214233), Abl (pET23a-H6-TEV-cAbl; AddGene: 214234), and AncSZ (pET23a-H6-TEV-AncSZ; AddGene 214235) were used to express and purify the four kinase domains, same as in reported before^33^. In short *E. coli* BL21(DE3) cells with the YopH tyrosine phosphatase-expressing helper plasmid were transformed with the constructs, and plated on ampicillin and streptomycin LB-agar plates. A single colony was used to grow an overnight starter culture that was used to inoculate 6 L of TB medium. The expression was induced by 0.5 mM IPTG at OD_600_ of 0.5 for 16-18 hours at 18 °C. The cells were harvested, resuspended in lysis buffer (50 mM Tris, pH 8.0, 300 mM NaCl, 20 mM imidazole, 10% glycerol), and captured on HisTrap HP column. The kinases were eluted from the HisTrap column with high imidazole directly onto on 5 mL HiTrap Q HP anion exchange column. Next, the kinases were eluted from the HiTrap Q HP column with an NaCl gradient. The AncSZ kinase was then run in a final step on a HiLoad 16/600 Superdex 75 pg column (Cytiva) using the size exclusion and storage buffer (10mM HEPES, pH 7.5, 100mM NaCl, 1 mM TCEP, 5 mM MgCl2 10% glycerol). For the other three kinases, the His_6_-tag was then cleaved using TEV protease, and the cleaved kinase domains were isolated after flowthrough through a HisTrap HP column. The kinases were then purified by size exclusion chromatography on a HiLoad 16/600 Superdex 75 pg column (Cytiva) using the size exclusion and storage buffer. Purified kinases were aliquoted, snap-frozen in liquid N_2_ and stored at –80 °C.

### Cloning of the X11 library

The X_11_ and X_8_ libraries used in this investigation were prepared similarly to the already available X_5_YX_5_ library reported previously^33^. In all cases the library is cloned at the N-terminus of the eCPX that contains a Myc-tag et the C-terminus. For this purpose, the eCPX-cMyc was PCR-amplified from the reported pBAD33-X_5_YX_5_-eCPX-cMyc-tag plasmid library using the link-eCPX-fwd and eCPX-rev primer. NNS-encoded oligonucleotide library with the 5’ SfiI restriction site and 3’ eCPX-linker overlapping sequence obtained from MilliporeSigma, eCPX-rand-lib-X11 was used as forward primer in combination with the eCPX-rev reverse primer to produce the X_11_-eCPX-cMyc-tag DNA library by PCR amplification. In parallel, the pBAD33 plasmid backbone was PCR-amplified using the eCPX-BB-for and eCPX-BB-rev. The pBAD33 PCR-amplified plasmid backbone was also treated with DpnI (NEB). Both the pBAD33 plasmid backbone and the eCPX-appended libraries were treated with the SfiI DNA restriction enzyme (NEB) after PCR-purification. The SfiI treated pBAD33 plasmid backbone was additionally treated with Quick CIP (NEB) as well. Both the DNA libraries and the pBAD33 plasmid backbone were then gel-purified and used for a ligation reaction in 5:1 molar ratio using the T4 DNA ligase (NEB). After PCR-purification of the ligation, the DNA was used to transform NEB 10-beta electrocompetent *E. coli* (NEB) using the Bio-Rad xCell electroporator (Bio-Rad) at 2kV, 25µF, and 200Ω. After a 1 hour recovery at 37 °C in 1 mL of the recovery medium supplied with the electro-competent *E*.*coli*, the transformed cells were used to inoculate 200 mL of LB containing 25 µg/mL chloramphenicol and grow a midi-preparation culture over-night at 37 °C and 215 rpm. The pBAD33-X_11_-eCPX-cMyc and the pBAD33-X_8_-eCPX-Myc plasmid libraries were then midi-prepped from the over-night cultures (ZymoPURE™ II Plasmid Midiprep Kit).

### Preparation of cells for display of phosphorylated peptide libraries

Bacterial display and phosphorylation of the displayed peptides was performed similarly to^33^, with small adjustments. In short, 50-100 ng of the X_5_YX_5_, X_11_ or X_8_ plasmid library was used to transform 25 µL of electrocompetent MC1061 F^-^ *E*.*coli* (Lucigen ≥4 × 10^10^ cfu/µg) in a 1 mm electroporation cuvette and micropulser (Bio-Rad) set at 1.8 kV. The transformed cells were recovered for one hour at 37 °C and 215 rpm in 975 µL of warm recovery medium supplied with the electrocompetent cells. For the display culture, 100 mL of LB containing 25 µg/mL chloramphenicol were inoculated with 0.98 mL of the transformed cells after recovery and grown at 37 °C and 215 rpm until reaching OD_600_ of 0.5-0.6. At this point 20 mL of the culture were induced with 0.4% (w/v) arabinose for 4 hours (+/- 30 minutes) at 25 °C and 215 rpm. Cells from 5 mL of the induced culture were collected by centrifugation at 4000 g, resuspended in 5 mL of PBS, and separated into several 750 µL aliquots. The cells of each aliquot were collected by centrifugation (4000 g), the supernatant was discarded, and cells were stored overnight at 4 °C. For peptide phosphorylation each cell pellet from a 750 µL aliquot was resuspended in 500 µL of kinase buffer pH 7.5, 50mM Tris, 10mM MgCl2, 150mM NaCl, 2mM Orthovanadate and 1 mM TCEP) and the volume of cells needed for the given experiments was used for phosphorylation of the displayed peptides by adding purified kinase domains c-Src, c-Abl, EPHB1 and AncSZ up to 2.5 µM, 5 mM of creatine phosphate, 50 μg/mL creatine phosphokinase (from rabbit muscle, Sigma), and 1 mM ATP. The phosphorylation reaction was incubated at 37 °C for 3 hours and stopped by adding EDTA up to 25 mM. The cells of the phosphorylated bacterial display library were then collected by centrifugation (4000 g) and resuspended in the same volume of the buffer used for the selection.

### Multi-round selection against SH2 domains

To perform the single selection experiment using the phosphorylated peptide library against, 75 µL of streptavidin coated magnetic beads (Dynabeads™ FlowComp™ Flexi Kit, Thermo-Fisher) were washed twice in 1 mL of SH2 binding buffer (50 mM HEPES pH 7.5, 150 mM NaCl, 0.05% Tween 20, 1mM TCEP) and incubated in total of 150 µL SH2 binding buffer containing 20 µM biotinylated SH2 domain on a rotator and 4 °C for 2-3 hours in low protein binding microcentrifuge tubes (1.5 mL, Thermo Scientific™). The functionalized magnetic beads are then washed twice in 1 mL of SH2 binding buffer, resuspended in 75 µL of the SH2 binding buffer and mixed with phosphorylated peptide-displaying cells resuspended in 100 µL of the same buffer containing 0.05% Tween 20. After incubation for one hour on a rotator and 4 °C in the low protein binding microcentrifuge tubes (1.5 mL, Thermo Scientific™), the supernatant containing the non-bound library fraction was discarded and the magnetic beads with the bound fraction were washed in 1 mL of the SH2 binding buffer containing Tween 20 for 30 minutes on a rotator and 4 °C. At this point the magnetic beads with the bound fraction were collected, resuspended in 100 µL of MilliQ water and incubated at 100 °C for 10 minutes to extract the plasmid DNA. As input library, 100 µL of the surface displayed library before phosphorylation was washed with MilliQ water through centrifugation, and the DNA was extracted same as the selection sample. The extracted DNA was then used both for deep sequencing sample preparation and for PCR amplification using the 3’SfiIX11-fwd and eCPX-rev primers to clone the enriched library in the pBAD33 plasmid backbone, as in the original libraries. The purified ligation was directly used for cell transformation and production of peptide displaying cells for the next round of selection.

### Display and phosphorylation quality control

For every set of experiments a new batch of phosphorylated peptide-displaying cells was produced. A sample from each batch of these phosphorylated peptide-displaying cells was used to control the quality and consistency by selection using an antibody that specifically binds to phosphorylated tyrosine, irrespective of the flanking sequence context^33^. For this quality control, 50 µL of the bacterial peptide library after phosphorylation were pelleted by centrifugation at 3000 g and resuspended in 50 µL PBS-BSA buffer (PBS + 0.2% BSA) containing 1:1000 Platinum *α*-PhosTyr 4G10 biotin conjugated antibody (Sigma). The cells were incubated with the antibody on ice for one hour and then pelleted through centrifugation at 3000 g to remove the non-bound antibody. In the same manner the cells were washed once in 100 µL in PBS-BSA, resuspended in 50 µL PBS-BSA, and mixed with 37.5 µL of streptavidin coated magnetic beads (Dynabeads™ FlowComp™ Flexi Kit, Thermo-Fisher) previously washed and resuspended in PBS-BSA. The cell-bead suspension was incubated on a rotator at 4 °C for 20 minutes to capture the library fraction labelled with the biotinylated antibody. After removing the non-bound fraction via magnetic separation, followed by a 15 minute wash in 0.5 mL in PBS-BSA, the plasmid DNA of bead-bound fraction of the surface-displayed library was extracted by incubation at 100 °C for 10 minutes in 50 µL of MilliQ water. The sample was then analyzed by deep sequencing.

### Deep sequencing and preparation of selection samples

DNA extracted from both the collected input and selection-enriched samples was analysed by deep sequencing. Each sample was prepared for sequencing similarly as reported before^33^. In short, the DNA samples were first PCR-amplified over 15 cycles using the Q5 polymerase master mix (NEB) and the primers altTruSeq-eCPX-fwd and altTruSeq-eCPX-rev. After analyzing small amounts of these PCR reactions on DNA-agarose gel, the PCR products were used for the second PCR amplification without any purification and roughly amount adjusted to the intensity of the bands in the control DNA gel, in order to normalize the input DNA levels. Sample-specific indexing primer pairs (from the D500 and D700 primer series, see **Table S7**) were used in this second PCR to label the samples. The second-round PCR products were then gel-purified and quantified on NanoDrop so that equal amounts of each sample were mixed together. The final sample mix, constituted to yield around 1 million reads for each sample, was quantified by qPCR using the NEBNext Library quantification kit (NEB) and the StepOnePlus cycler (Applied Biosystems) to produce the needed DNA concentration of 4 nM. For deep sequencing on the Illumina MiSeq system using the MiSeq Reagent Kit v3 (150 cycle) the sample was denatured in 0.1 M NaOH for 5 minutes at room temperature and diluted down to 20pM. The quality of each sequencing run was controlled by adding up to 5% of denatured and diluted down to 20pM PhiX (Illumina). Demultiplexing of the data by sample using the defined index pairs was performed automatically by Illumina BaseSpace.

### Measuring affinities of SH2 domains and phosphopeptides using competitive fluorescence polarization

Measured binding constants (*K*_D_) from competition mode fluorescence polarization experiments were obtained using the purified SH2 domains used for the selection and chosen sets of peptides with an phosphorylated tyrosine exactly as described previously^33^. For all Src-family kinase-derived SH2 domains, the fluorescently labelled peptide FITC-Ahx-GDG(pY)EEISPLLL (c-Src SH2 consensus) was used as the probe, and for the Grb2 SH2 domain, the fluorescently labelled peptide FITC-Ahx-FDDPS(pY)VNVQN (Ahx = 6-aminohexanoic acid) from the Y177 phospho-tyrosine site of BCR. Before performing the competition mode fluorescence polarization measurement, the *K*_D_ of the probes for the respective SH2 domains was first measured in a direct mode of fluorescence polarization. The sets of non-labelled peptides-sequences for the competition-based measurement were chosen in different ways: For the Grb2 and Fyn SH2 domains, the choice was based on sequences found during selections against these SH2 domains, so that they span a larger range of predicted relative affinities. For the set of peptides where single point mutations were compared, phosphopeptides reported in the PhosphoSitePlus database and previously shown to bind to the c-Src SH2 domain^33^, were chosen alongside their respective single point mutants. To choose phosphopeptides for validation of the Lyn SH2 domain, we started from all verified phosphopeptides found in the PhosphoSitePlus database, filtered based on their co-expression with the Lyn kinase using the STRING database. Using the validation data for the Fyn SH2 domain, the range of predicted relative binding affinities that corresponds to the range between 1 nM and 100 µM, which is the dynamic range of reliable measurement of absolute binding affinities using our competitive fluorescence polarization setup, was defined and used to further filter the set of candidate phosphopeptides. A subset that evenly covers the chosen range of predicted relative affinities was finally picked for the validation measurements of the Lyn SH2 domain. For the c-Src, Yes, and Blk SH2 domains, a combination of the previously chosen peptides were used, after verifying that their predicted relative affinities for the given SH2 domain fell within the same measurable range. All peptides containing phosphorylated tyrosine, both fluorescently labelled and non-labelled, were produced by SynPeptide (China).

### Processing of Sequencing Reads

For each sequencing library, the read pairs were first joined using FLASH-1.2.11 software package^47^, and the combined reads were scanned for a 33-bp region flanked by GTAGCTGGCCAGTCTGGCCAG to the left and GGAGGGCAGTCTGGGCAGTC right, allowing for up to five mismatches. After cropping out the flanking sequences, the remaining reads were filtered for exact matches to the library design, whereupon reads with a PHRED score below 20 at any position were discarded. The remaining reads were then translated to protein code. Count tables containing the number of occurrences of each sequence in the relevant input and bound libraries were finally created for each selection round of each experiment.

### Fitting Binding Models

To learn binding models based on sequencing data, ProBound minimizes a loss function consisting of a log-likelihood term and a regularization term:

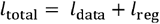

ProBound was thus configured to load one or more of count tables (one per selection round), model each table as a single-round SELEX experiment, and then compute the scaled binomial log-likelihood:

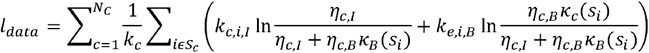

Here *N*_*c*_ is the number of count tables, *k*_*c*_ is the total number of reads in count table *c, k*_*c,i,I*_ and *k*_*c,i,B*_ are the number of reads observed for sequence *s*_*i*_ in the input and bound columns of count table *c*, and *η*_*c,I*_ and *η*_*c,B*_ are parameters controlling the predicted sequencing depth. κ_*c*_ (*S*_*i*_) is the predicted enrichment of sequence s_*i*_ in the bound library. ProBound was configured to model this enrichment using two ‘binding mode’ terms, one representing non-specific binding and experimental biases (*NS*) and one term representing sequence-specific SH2-domain binding (*S*):

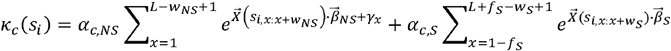

These terms are similar in that each receives additive contributions from all offsets along the sequence, and each contribution depends log-linearly on the subsequence starting at the offset in question. Mathematically, 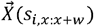 is a predictor vector that uses one-hot encoding to represent the subsequence *s*_*ix*_._*x+w*_ starting offset *x* and extending *w* residues to the right. The coefficient vectors (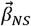 and 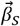) encode the log-fold effects that the residues in the substring have one the binding (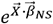 and 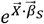). The components in 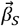 thus correspond to the entries in the free-energy matrix —ΔΔ*G*/*RT* that we wish to learn. The overall scale of the selection was set by the ‘activity’ weights *α*_*c,NS*_ and *α*_*c,s*_, which took independent values for each count table *c*.

Beyond the similarities discussed above, the binding models were configured to differ in important ways. For the non-specific mode to focus on simple sequence dependences, short subsequences of length *w*_*NS*_ =3 were used (for Grb2, *w*_*NS*_ =1 was used since longer substrings allowed this mode to capture the main motif YXNX. This was also done for the pTyrVar library). A position-specific bias term γ*x* was also included to let this mode absorb non-homogenous sequence biases. In contrast, the sequence-specific binding mode was focused on extended SH2 binding sites by using substring of length *w*_*S*_ = 11 and by not including a position-specific bias term. Moreover, because the variable region in the library (which had length *L* = 11) was flanked by the constant sequences GQSGQ on the left and GGQSG on the right, and because these sequences in principle could be a part of a binding site, the sum over offsets *x* was extened to include substrings overlapping up to *f*_*S*_ = 5 flanking residues. Finally, to focus this mode on phosphotyrosine-specific binding, the coefficient vector 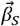 was constrained to assign weight zero to a tyrosine at the central position and weight -10 for all other residues. The regularization term *l*_*reg*_ consisted of a *L*_2_regularizer, an exponential barrier term, and a Dirichlet term (with count 5) as described in the original publication^34^.

## Data and software availability

The raw sequencing data, processed count tables, configuration files, and two supplemental tables are available at bussemakerlab.org/papers/SH2/.

ProBound source code is available at github.com/RubeGroup/ProBound.

## ACKNOWLEDGEMENTS

We thank the members of the Shah and Bussemaker labs for valuable discussions. This work was supported by National Institute for Mental Health Research Grant R01MH106842 to H.J.B., National Institute for General Medical Sciences Research Grant R35GM138014 and American Cancer Society grant RSG-23-1038049-01-TBE to N.H.S., as well as National Cancer Institute Cancer Center Support Grant P30CA013696 and National Center for Advancing Translational Sciences Grant UL1TR001873. The content is solely the responsibility of the authors and does not necessarily represent the official views of the NIH.

## AUTHOR CONTRIBUTIONS

DG, HTR, NHS, and HJB designed research; HTR, CR, LANM, and XL analyzed data; DG and RV performed experiments; NHS performed structural analyses; DG, HTR, NHS, and HJB wrote the manuscript; all authors reviewed and approved the manuscript.

## LIST OF SUPPLEMENTAL TABLES

**Table S1:** Next-generation sequencing datasets generated.

**Table S2:** List of ProBound analyses, settings, and resulting binding models.

**Table S3:** Validation measurements, unpaired.

**Table S4:** Predictions for peptides in the PhosphoSitePlus database, for cSrc, Fyn, Lyn, Grb2.

**Table S5:** Validation measurements, single-amino-acid substitution pairs.

**Table S6:** Predictions for peptides in the PTMVar database, for cSrc, Fyn, Lyn, Grb2.

**Table S7:** Cloning and sequencing primers used.

## SUPPLEMENTAL FIGURES

**Figure S1:**
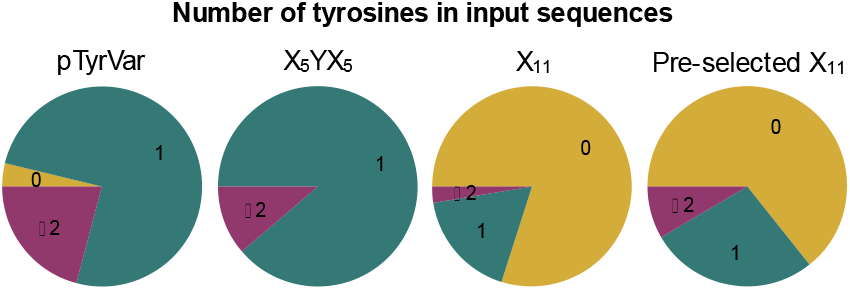
Proportion of sequences containing zero, one, or two or more tyrosine residues in the different input libraries.

**Figure S2:**
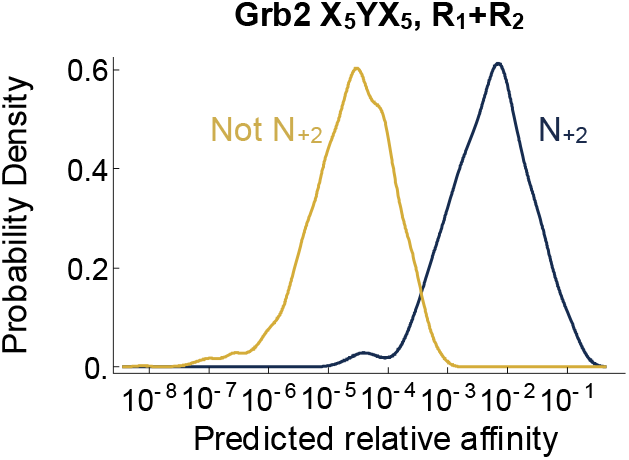
Impact of N_+2_ on predicted Grb2 binding. Plot shows the distribution of binding affinities (shown using a log-scale kernel density estimator) predicted by the Grb2 model in from Fig. 4a. Sequences containing an N_+2_ (blue) are grouped separately from the other sequences (orange).

**Figure S3:**
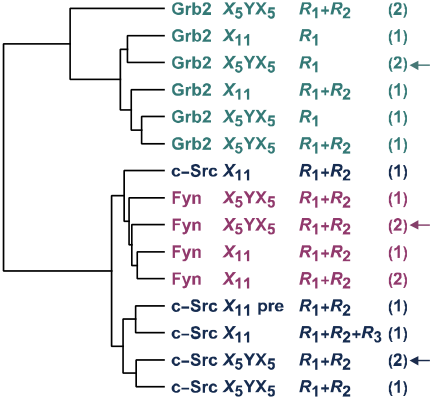
Broader comparison of multi-round models for multiple SH2 domains. The dendrogram shows the clustering of various binding models for the c-Src, Grb2, and Fyn SH2 domains, built using ProBound from data generated using different starting libraries, number of selection rounds. Numbers in parentheses denote replicates. Arrows denote the X_5_YX_5_ models used for all other analyses in this paper.

**Figure S4:**
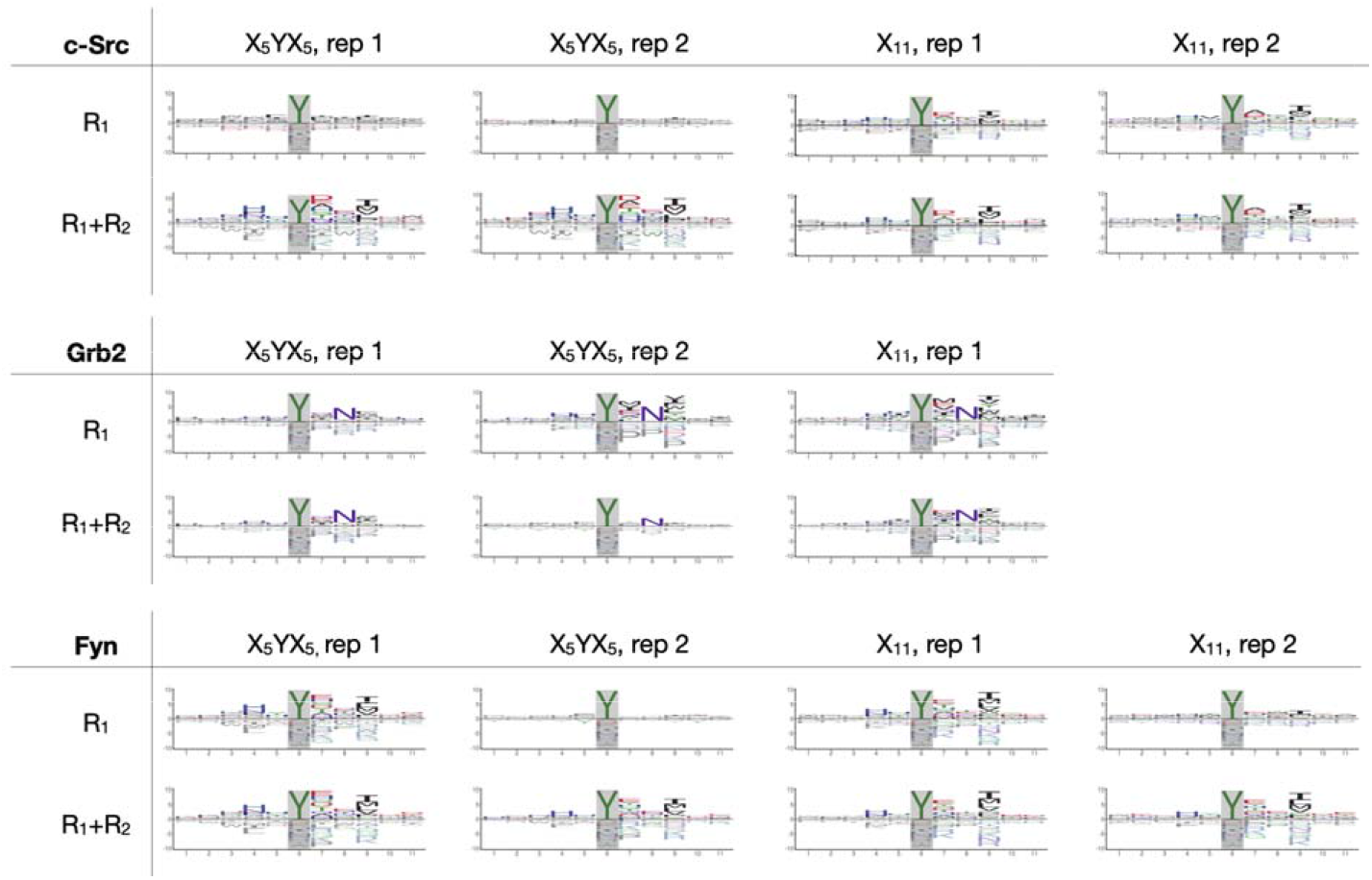
Binding models for the c-Src, Grb2 and Fyn SH2 domains. The models were learned using different combinations of starting libraries (X_5_YX_5_ or X_11_), selection round (R_1_ or R_1_+R_2_), and replicates (rep 1 or rep 2).

**Figure S5:**
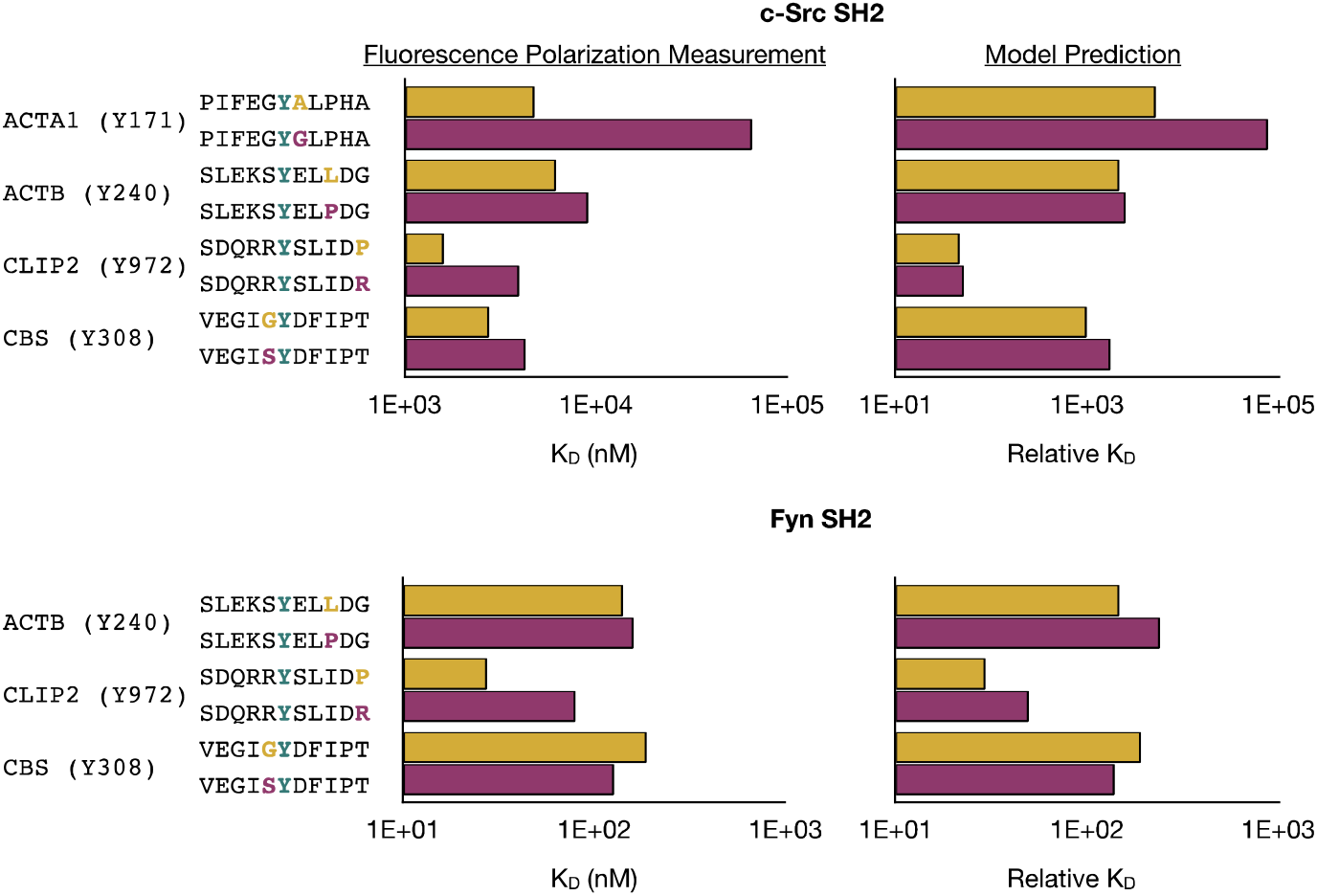
Impact of single-amino-acid substitutions on c-Src and Fyn SH2 binding. The bar charts show the measured K_D_ value (left) and the predicted relative K_D_ (right) for pairs of naturally occurring sequence variants (highlighted letters). The predictions were made using the models shown in Fig. 4a.

